# Inhibition and Disassembly of Tau Aggregates by Engineered Graphene Quantum Dots

**DOI:** 10.1101/2022.12.29.522245

**Authors:** Runyao Zhu, Kamlesh M. Makwana, Youwen Zhang, Benjamin H. Rajewski, Juan R. Del Valle, Yichun Wang

**Affiliations:** Department of Chemical & Biomolecular Engineering, University of Notre Dame, Indiana 46556, United States; Department of Chemistry & Biochemistry, University of Notre Dame, Indiana 46556, United States

## Abstract

Tauopathies are a class of neurodegenerative diseases resulting in cognitive dysfunction, executive dysfunction, and motor disturbance. The primary pathological feature of tauopathies is the presence of neurofibrillary tangles in the brain composed of tau protein aggregates. Although numerous small molecules are known to inhibit tau aggregation, it is still challenging to use them for therapeutic applications due to their limitations in specific targeting and the blood-brain barrier (BBB) penetration. Graphene quantum dots (GQDs), one of graphene nanoparticles, can penetrate the BBB and are amenable to functionalization for targeted delivery. Moreover, these nanoscale biomimetic particles can self-assemble or assemble with various biomolecules including proteins. In this paper, for the first time, we showed that GQDs interacted with tau proteins *via* electrostatic and π-π stacking interactions to inhibit the fibrillization of monomeric tau and to trigger the disaggregation of tau filaments. *In vitro* thioflavin T assays demonstrated that negatively charged GQDs with larger sizes inhibited tau aggregation more efficiently, while positively charged ones were more effective in the disassembly of tau fibrils. Moreover, GQDs blocked the seeding activity of tau fibrils in a cellular propagation assay. Overall, our studies indicate GQDs with engineered properties can efficiently inhibit and disassemble pathological aggregation of tau proteins, which supports their future developments as a potential treatment for tauopathies.

## Introduction

Pathogenic aggregation of tau proteins into neurofibrillary tangles (NFTs) is a characteristic feature of tauopathies, which are a group of neurodegenerative diseases, including Alzheimer’s disease (AD),^1^ frontotemporal dementias (FTDP-17),^2^ Pick’s disease,^3^ and progressive supranuclear palsy.^4^ The normal function of tau is to stabilize microtubules by binding with tubulin in neurons. These microtubules provide a structural backbone for axons and dendrites and serve as ‘railways’ for the transport of proteins and organelles in axons and dendrites.^5^ Pathogenic fibrillization of tau reduces cytoskeletal stability, interferes with synaptic transmission and axonal transport, and causes aberrant communication between neurons.^6^ These insoluble tau deposits can arise due to gene mutations or aberrant post-translational modifications such as hyperphosphorylation and acetylation.^2,7,8^ In addition, tau aggregates can spread from neuron to neuron and induce the misfolding of normal soluble tau, leading to the propagation of tau pathology in a prion-like manner.^9,10^ Small molecule library screening campaigns have identified several compounds that either directly or indirectly inhibit tau aggregation.^11,12^ These molecules include modulators of tau post-translational modification,^8^ microtubule stabilizers,^13^ tau assembly inhibitors,^13,14^ and degradation promoters.^15^ However, there are no disease-modifying drugs that target tau aggregation. Moreover, many drug candidates suffer from limited penetration of the blood-brain barrier (BBB) and off-target effects, reducing central nervous system bioavailability and limit their clinical utility.^15,16^ Furthermore, compounds targeting kinases upstream of tau may disrupt other physiological processes resulting in undesirable side effects including cancers.^8^ Therefore, developing novel drugs to overcome current challenges is critical for the treatment of neurodegenerative tauopathies.

Nanoparticles (NPs), as drug nanocarriers, have demonstrated the ability to deliver drugs into the brain and are capable of controlled release and targeted delivery.^17,18^ Graphene quantum dots (GQDs), are single or a-few layered graphene with lateral dimensions typically less than 20 nm.^19^ These nanosheets have garnered much attention in biomedicine, including bioimaging, biosensor, and cancer phototherapy, due to their unique optical, electrochemical, and physicochemical properties, high biocompatibility, and low cytotoxicity.^20-23^ Moreover, GQDs can be used as nanocarriers in drug delivery systems upon conjugation with target ligands and chemotherapeutic agents *via* either covalent or noncovalent bonds because of their surface functional groups and π-conjugated planar structure.^24^ GQD-based drug delivery systems can penetrate the BBB and specifically target brain tumor regions, unraveling great potential in neuroscience therapeutics.^25-27^ Most importantly, GQDs exhibit a wide spectrum of biomimetic properties, including polarizability, amphiphilic character, and participation in electrostatic interaction, hydrophobic interaction, hydrogen bonding, and π-π stacking.^28,29^ Thus, GQDs can self-assemble ^30^ or assemble with various proteins, such as amyloid-β (Aβ) peptides,^31,32^ biofilm amyloids,^28^ and α-synuclein,^25^ demonstrating their potential to protect against the neuronal damage associated with AD and Parkinson’s disease. Furthermore, the properties of GQDs, such as size,^33^ charge,^34^ functional groups,^35^ and chirality,^36^ can be engineered precisely, affording opportunities to modulate their assembly with proteins.^37,38^

Here, we report the first example of inhibition of tau aggregation and disassembly of tau filaments by engineered GQDs. To investigate the effect of GQDs on tau aggregation, we engineered GQDs with different sizes and charges. By monitoring the amount of tau aggregates, the secondary structure, and the morphology of tau, we found that negatively charged GQDs with a larger size inhibited tau aggregation more effectively. In contrast, GQDs with positive charge disassembled more tau mature fibers than GQDs with negative charge. These findings were potentially attributed to the fact that aromatic residues on the aggregation-prone region of tau can interact with GQDs *via* π–π stacking interactions verified by the fluorescence quenching assay. In addition, the positively charged microtubule-binding domain of tau can interact with carboxyl groups on the edge of GQDs *via* electrostatic interactions proved by thioflavin T (ThT) aggregation assay results. Cellular propagation assay to test the cellular transmission of tau fibers demonstrate that GQDs can block the aggregation of endogenous tau induced by extracellular seeds. Overall, our findings suggest that engineered GQDs are a promising and effective platform for the development of therapeutics targeting tau propagation.

## Results and discussion

As-synthesized GQDs carrying carboxyl groups on the edges in an average diameter of 8 nm were obtained by the previously reported method **(Fig. 1a, b, c, Fig. S1)**.^33^ The zeta potential of as-synthesized GQDs was -19.9 ± 2.30 mV **(Fig. 1d)**. To obtain GQDs with different charges, we reduced the negative charge by functionalization of the GQDs with cysteine (Cys) and ethylenediamine (EDA) *via* carboxyl groups **(Fig. 1a, c)**. The zeta potentials of Cys-GQDs and EDA-GQDs were -1.46 ± 0.42 mV and 1.63 ± 0.55 mV, respectively, **(Fig. 1d)** due to the pI value of Cys, 5.02, and pKa value of EDA, 10.7.

**Fig. 1.**
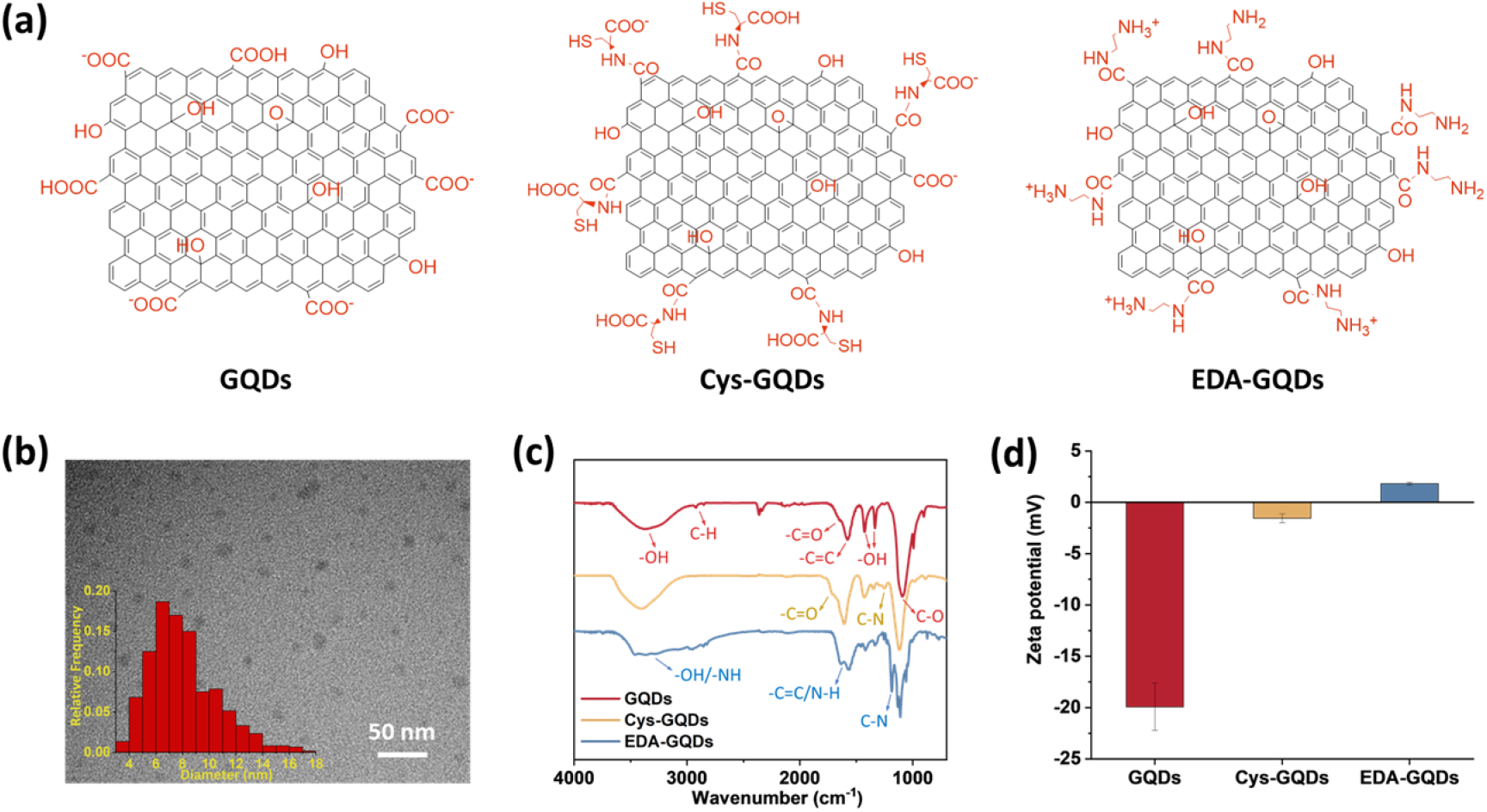
(a) Molecular structure, (b) Transmission electron microscope (TEM) image and size distribution of GQDs. The average diameter is 8.1 ± 2.5 nm. (Scale bar: 50 nm) (c) Fourier transform infrared (FTIR) spectra and (d) zeta potentials of GQDs, Cys-GQDs, and EDA-GQDs.

Following the synthesis, characterization, and functionalization of GQDs, we investigated the charge effect of GQDs on the aggregation of a recombinant 0N4R isoform of tau. Tau used in these studies includes a P301L mutation (tau_P301L_) which is frequently observed in patients with FTDP-17 **(Fig. S2)**.^39^ This missense mutation leads to a two-fold increase in aggregation rate compared with the full-length tau.^40^ Tau_P301L_ contains four microtubule-binding repeat domains (R1-R4) that harbor two key aggregation-prone motifs: PHF6* (VQIINK, residues 217-222) and PHF6 (VQIVYK, residues 248-253)._7_ A large number of basic residues result in an overall positive charge in the repeat domains at physiological pH **(Fig. 2a)**. Firstly, the aggregation of tau_P301L_ was monitored by a ThT fluorescence assay **(Fig. 2b)**. ThT is a benzothiazole dye that exhibits enhanced fluorescence upon binding to tau fibers with β-sheet secondary structure.^41^ In the presence of heparin sulfate, tau_P301L_ was induced to aggregate, resulting in a sigmoidal curve from increased ThT fluorescence **(Fig. 2c)**. Incubation of tau_P301L_ in the presence of heparin and 6.25 μM GQDs or Cys-GQDs resulted in 99% inhibition of endpoint ThT fluorescence **(Fig. 2c, d)**. However, EDA-GQDs with overall positive charge exhibited 88.8% inhibitory activity, less effective than the negatively charged GQDs and Cys-GQDs.

**Fig. 2.**
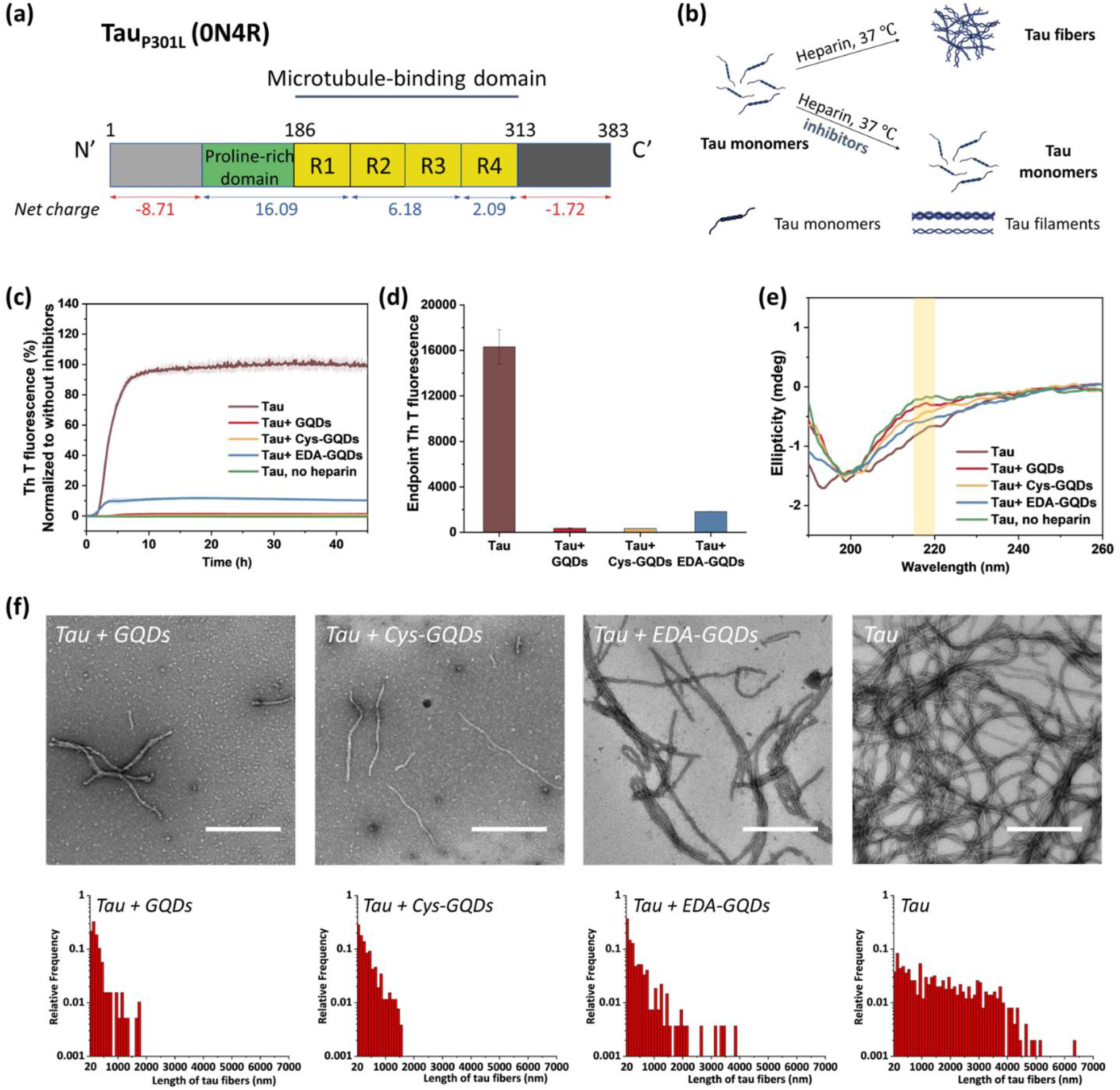
Tau_P301L_ aggregation was inhibited by GQDs, Cys-GQDs, and EDA-GQDs. (a) Aggregation-prone tau_P301L_ sequence. (b) Schematic representation of the *in vitro* aggregation of recombinant tau (10 μM) in the presence and absence of inhibitors at a concentration of 6.25 μM. (c-d) In the presence of GQDs or functionalized GQDs, thioflavin T (ThT) fluorescence of tau aggregation was reduced significantly. The inhibitory effect of EDA-GQDs is weaker than GQDs and Cys-GQDs. (e) Circular dichroism (CD) spectra, (f) TEM images, and length distributions of tau protein incubated with/without 6.25 μM GQDs or functionalized GQDs for 4 days. (Scale bars: 500 nm)

The mechanism of tau aggregation in the presence of heparin is widely described as ligand-induced nucleation-dependent polymerization (NDP).^42^ The lag time (***t***_***lag***_) and the apparent elongation rate constant (***k***_***app***_) of the aggregation process were derived by fitting each kinetic data to a Gompertz growth function **(Table S1)**.^43, 44^ The lag time represents the time of nucleation process (i.e., initial seed formation of tau) and the apparent elongation rate constant represents the rate of growth process (i.e., the elongation of tau fibrils). The tau aggregation without GQDs as a control showed a ***t***_***lag***_ of 1.84 ± 0.01 h and a ***k***_***app***_ of 0.76 ± 0.008 h^-1^. In the presence of GQDs or Cys-GQDs, ***t***_***lag***_ was increased to nearly double that of the control sample, while ***k***_***app***_ was reduced to 0.70 ± 0.039 and 0.38 ± 0.021 h^-1^, a decrease of 7.9% and 50 % compared to the control sample, respectively. These results suggest that GQDs and Cys-GQDs impeded both seed formation and the fibril elongation processes. In contrast, tau incubated with EDA-GQDs led to a significant decrease in ***t***_***lag***_ while ***k***_***app***_ slightly increased. This is potentially caused by the presence of EDA-GQDs that increase the local concentration of soluble monomeric tau and thus facilitate the initial formation of aggregation-prone tau seeds. Despite a decreased ***t***_***lag***_, EDA-GQDs were able to reduce the formation of tau fibers by 88.8 % after 5 h incubation compared to the control sample. We detected the effect of GQDs and functionalized GQDs on the secondary structure transition of tau_P301L_ using circular dichroism (CD) spectroscopy **(Fig. 2e)**. In the absence of heparin, soluble tau monomers showed a negative peak at ∼200 nm, indicative of a random coil conformation. After heparin-induced fibrillization, a growing CD signal at ∼218 nm, typical of β*-*sheet conformation, emerged.^45^ Upon incubation with GQDs, Cys-GQDs, or EDA-GQDs for 4 days, the CD intensity of tau at 218 nm was reduced by 61%, 39%, and 18% compared to the control sample, respectively, suggesting inhibition of transition to a β-rich tau assembly. The secondary structure of tau had less β-sheet content in the presence of GQDs or Cys-GQDs than in the presence of EDA-GQDs, which is consistent with the ThT assay results **(Fig. 2c)**.

The morphological features of tau_P301L_ fibrils were examined using transmission electron microscopy (TEM) after incubation for 4 days. To quantify the tau fibers in each sample, we measured their length and density using ImageJ **(Fig. 2f, Table S2)**. The aggregation of tau proteins in the control sample formed long filaments densely with the average length of 1623 ± 1255 μm and the average density of 4.53 ± 2.08 fibers/μm^2^. In the presence of GQDs or functionalized GQDs, heparin-induced tau_P301L_ assembled into shorter fibers with lower density. When tau_P301L_ monomers were incubated with GQDs, Cys-GQDs, or EDA-GQDs, the average length of formed fibers was decreased to 17%, 20%, and 35% of the control sample, while the average density of fibers was decreased to 30%, 34%, and 78%, compared to that of the control sample. Among samples incubated with the three GQDs, tau incubated with EDA-GQDs resulted in the formation of 3 to 4-fold more fibers per unit area than tau with GQDs or Cys-GQDs. The collective results indicate that GQDs and Cys-GQDs with negative charge have a better inhibitory effect on the fibrillization of tau than EDA-GQDs with positive charge.

To further investigate whether the amount of negative charges on GQDs can influence their inhibitory efficiency, we monitored the ThT fluorescence of tau_P301L_ incubated with GQDs or Cys-GQDs at different concentrations. GQDs and Cys-GQDs inhibited aggregation in a dose-dependent manner **(Fig. 3a)**. We obtained IC_50_ value for GQDs and Cys-GQDs by fitting the dose-response results to a sigmoidal function (**Fig. 3b)**. The IC_50_ of GQDs and Cys-GQDs are 0.70 μM, and 0.89 μM.^12,14^ These results suggest that when the zeta potential of GQDs is between -20 mV and -1.5 mV, the amount of negative charges on GQDs does not affect the inhibitory efficiency significantly.

**Fig. 3.**
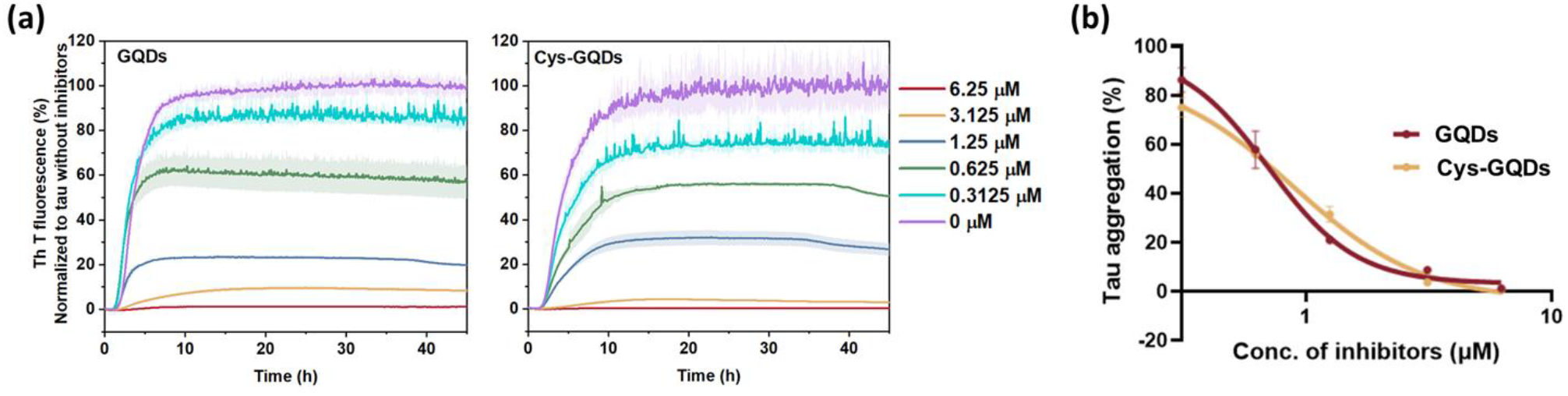
(a) ThT fluorescence of tau aggregation was decreased with the increased concentration of GQDs and Cys-GQDs. (b) Dose-response results of ThT fluorescence assay of GQDs and Cys-GQDs were fitted to a sigmoidal model. The IC_50_ of GQDs and Cys-GQDs is 0.70 μM and 0.89 μM, respectively, obtained from the dose-response curve.

One of the key characteristics of tau NFTs is cellular transmission from neuron to neuron, inducing the propagation of tau pathology.^9,10^ To examine whether GQDs and Cys-GQDs can inhibit the cellular seeding of recombinant tau, we employed HEK293 biosensor cells that stably expressed a tau-yellow fluorescent protein fusion [tau-RD(LM)-YFP] **(Fig. 4a)**.^46^ After incubating preformed tau_P301L_ fibrils with these cells for 48 h, aggregation of endogenous tau induced by extracellular seeds can be monitored by detection of focal puncta with green fluorescence. Pre-treatment of tau_P301L_ monomers with 0.12 μM of GQDs or Cys-GQDs in the presence of heparin decreased the intracellular aggregates by 60 % relative to the control treatment. As concentrations of GQDs and Cys-GQDs increased, the seeding activity of tau_P301L_ decreased. With 0.6 μM of GQDs or Cys-GQDs, the seeding activity of tau_P301L_ was almost completely abolished **(Fig. 4b)**. These results demonstrate that GQDs and Cys-GQDs prevent the fibrillization of tau monomers and thus block the cellular seeding activity of recombinant tau.

**Fig. 4.**
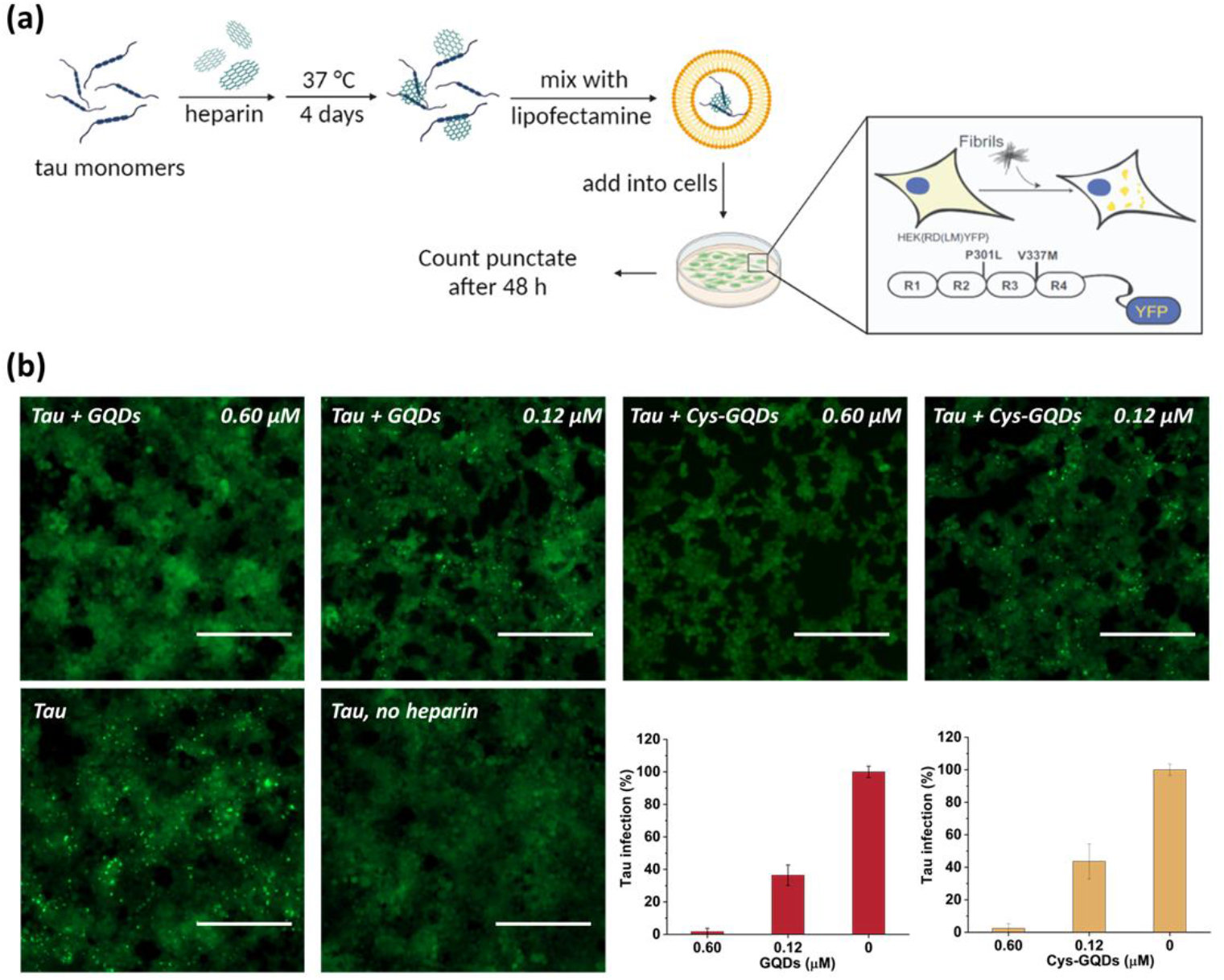
(a) Scheme of HEK293 cellular tau biosensor propagation assay. (b) GQDs and Cys-GQDs prevented the cellular seeding of tau_P301L_ monomers. Soluble monomeric tau_P301L_ (0.19 μM) in the presence of heparin was incubated with GQDs or Cys-GQDs for 4 days and then added to HEK293 cells stably expressing tau-RD (P301L/V337M)-YFP. Representative images of cells were taken at 20× magnification under fluorescein isothiocyanate (FITC) channel (ex: 469 nm/em: 525 nm). The green puncta with high fluorescence represented the aggregation of tau in cells induced by exogenous tau fibers. Scale bar: 200 μm. Bar graphs show the number of intracellular fluorescent puncta relative to control infection wells without inhibitors.

Tau NFTs are found within the neurons of patients with tauopathy, and thus it is important to investigate the effect of GQDs on the disaggregation of preformed tau fibers. Therefore, we performed the disaggregation of tau mature fibrils using ThT assay **(Fig. 5a)**. After incubation with GQDs, Cys-GQDs, or EDA-GQDs, the intensity of ThT fluorescence in the presence of tau fibrils decreased with time **(Fig. 5b)**, suggesting that tau fibrils were dissociated into tau monomers and oligomers. Based on endpoint ThT fluorescence after 30 h **(Fig. 5c)**, GQDs, Cys-GQDs, and EDA-GQDs induced 56%, 81%, and 92% disassembly of tau fibrils, which indicates that EDA-GQDs are more effective at disaggregating tau_P301L_ fibrils than negatively charged GQDs and Cys-GQDs. This is potentially due to the negatively charged fuzzy coat of tau_P301L_ fibrils comprised of the unfolded C-terminal and N-terminal domains **(Fig. 2a)** that surrounds the fibrillar tau core.^47^ The negative charges on the outermost layer of tau fibers can impede the interaction between the fibrillar tau core and negatively charged GQDs.

**Fig. 5.**
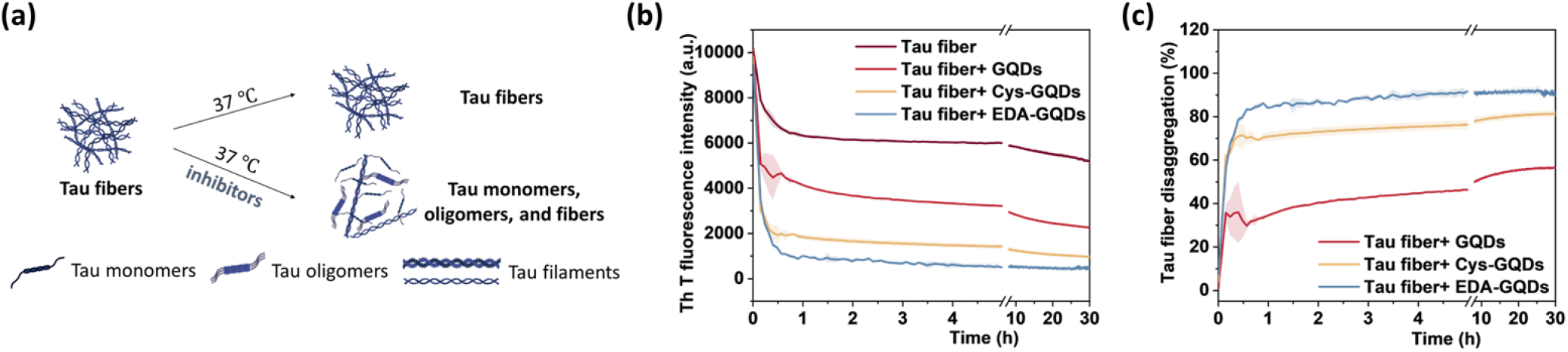
(a) Scheme of the *in-vitro* disaggregation assay of tau fibers (5 μM) in the presence and absence of inhibitors (6.25 μM). (b) ThT fluorescence of tau fibers and (c) the percentage of disaggregation of preformed tau fibers after incubation with GQDs, Cys-GQDs, or EDA-GQDs. The disassembly ability of GQDs and Cys-GQDs is weaker than EDA-GQDs.

To further confirm the disassembly ability of GQDs and Cys-GQDs, we tested if they can block the cellular transmission of mature tau fibrils in the tau biosensor seeding assay. After 36 h treatment of GQDs or Cys-GQDs with preformed tau fibrils, HEK293 cells expressing tau-RD(LM)-YFP were incubated with the pre-treated tau fibrils. With the increase in concentration from 0.12 μM to 0.5 μM, GQDs reduced the aggregation of endogenous tau from 18% to 80%, while Cys-GQDs reduced the intracellular aggregates from 6% to 88% compared to the control sample without GQDs or Cys-GQDs **(Fig. 6)**. These results suggest that GQDs and Cys-GQDs effectively block the seeding capacity of mature tau fibrils through disassembly into seed incompetent monomers. Lastly, to confirm that GQDs and Cys-GQDs are not toxic to neurons, we employed the CCK-8 assay with SH-SY5Y human neuroblastoma cells. GQDs and Cys-GQDs exhibited no appreciable toxicity toward SH-SY5Y cells up to 5 μM after 48 h **(Fig. S3)**.

**Fig. 6.**
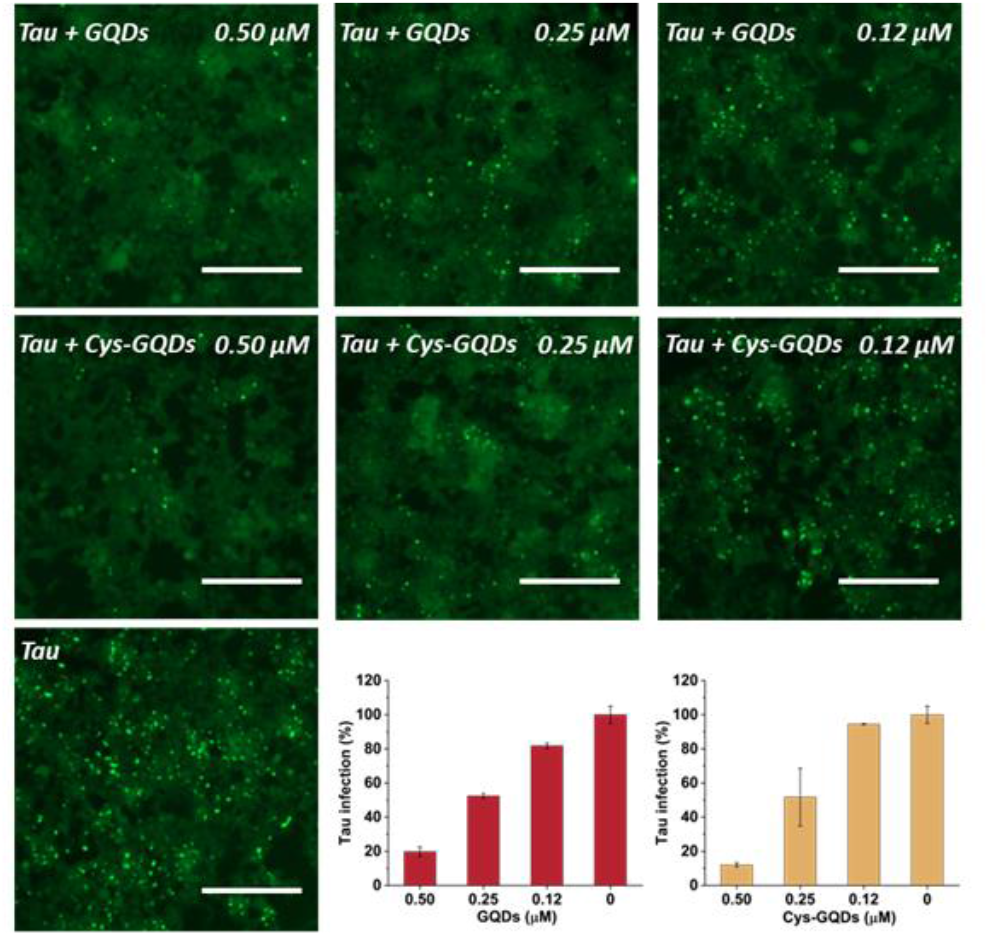
GQDs and Cys-GQDs prevented the cellular seeding of mature tau fibers. Preformed tau fibrils (0.19 μM) were treated with GQDs or Cys-GQDs for 36 h, then added to cells, and incubated for an additional 48 h. Representative images of cells were taken at 20× magnification under FITC channel (ex: 469 nm/em: 525 nm). The green puncta with high fluorescence represented the aggregation of tau in cells induced by exogenous tau fibers. Scale bar: 200 μm. Tau infection (%) in the bar graph shows the number of intracellular fluorescent puncta relative to control infection wells lacking the inhibitors.

Based on our experimental results, negatively charged GQDs can inhibit the fibrillization of tau proteins more efficiently than GQDs with positive charge. This suggests that the positively charged domains of tau, especially the aggregation-prone R2 and R3 domains, prefer to engage in more electrostatic interactions with the carboxyl groups on the edge of GQDs rather than form hydrogen bonds that are critical for the formation of tau aggregates.

Moreover, based on ThT aggregation assay results **(Fig. 2c)**, there are other factors in addition to the negative charge of GQDs that may result in the inhibition of tau aggregation. We investigated the other potential interactions between tau and GQDs using fluorescence spectroscopy. Generally, the tau exhibits intrinsic fluorescence emission near 302 nm when excited at 265 nm, resulted from tyrosine (Tyr) fluorophores in the tau structure.^48,49^ When 10 μM of tau was mixed with GQDs or Cys-GQDs in increasing concentrations, the fluorescence intensity of tau was significantly quenched **(Fig. 7)**, suggesting that the π-conjugated planar structure of GQDs binds with Tyr residues in tau, probably *via* π-π stacking interactions.

**Fig. 7.**
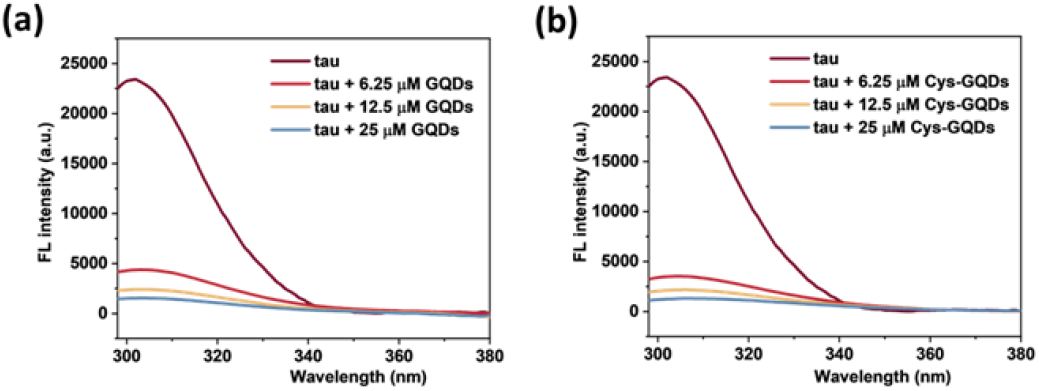
Fluorescence spectra of tau protein (10 μM) incubated with 6.25, 12.5, and 25 μM of (a) GQDs and (b) Cys-GQDs (Ex: 265 nm). The fluorescence of tau was quenched significantly in the presence of GQDs or Cys-GQDs.

Besides the effect of charge, we also found that the inhibitory efficiency of GQDs increased significantly as the diameter of GQDs increased from 6.5 nm to 13.5 nm based on ThT fluorescence assay **(Fig. S4)**. The diameter of the full-length tau monomer is 13 ± 0.6 nm.^50^ As the diameter of GQDs increases from 6.5 nm to 13.5 nm, single GQDs with a larger diameter are able to interact with more tau monomers on the edge *via* electrostatic interaction than the ones with a smaller diameter. Also, when interacting with larger GQDs, the spacing between two tau monomers increases, reducing the local concentration of tau and enhancing inhibitory efficiency. In addition, increasing the size of GQDs promotes more aromatic residues in tau to bind to the surface of GQDs *via* π-π stacking, thereby increasing the binding strength between tau and GQDs.

Overall, the potential mechanism of the interaction between GQDs and tau protein can be summarized in **Fig. 8**. For the aggregation of tau_P301L_ (**Fig. 8a)**, negatively charged GQDs with carboxyl groups can interact with repeat domains of tau with positive charge *via* electrostatic interactions. Meanwhile, GQDs with π-conjugated planar structure can interact with amino acids containing aromatic rings in tau proteins *via* π-π stacking, including histidine, phenylalanine, and tyrosine. The carbon-conjugated plane of GQDs could also interact with positively charged amino acids in tau *via* cation-π interactions. For the disassembly of tau fibrils **(Fig. 8b)**, GQDs with negative charge exhibited reduced efficacy, which may be due to the negatively charged fuzzy coat surrounding the fibril core of tau.

**Fig. 8.**
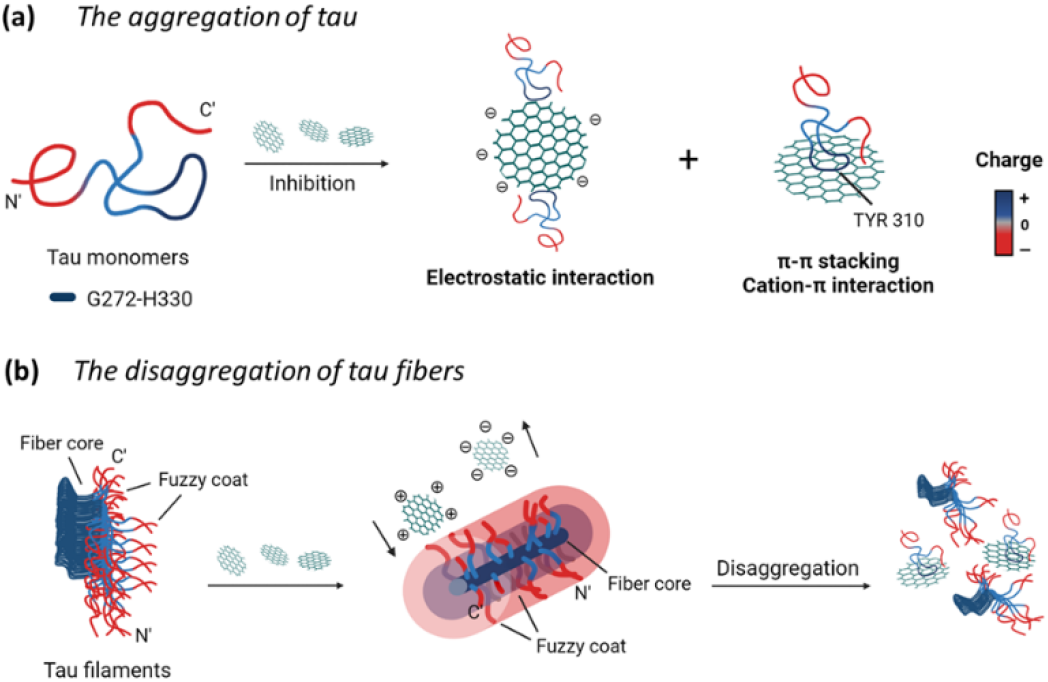
Schematics of the charge effect of GQDs on their interactions with tau proteins. GQDs can interact with tau proteins *via* electrostatics interactions and π-π stacking, thereby (a) preventing the tau aggregation, and (b) inducing the disassembly of tau fibers.

## Experimental

### Synthesis and characterization of GQDs

GQDs were prepared using a modified Hummers method.^33^ 0.45 g of carbon fiber was added into 90 mL of concentrated H_2_SO_4_ (98%) and stirred for 1.5 h. After stirring, 30 mL of concentrated HNO_3_ (68%) was added into the above mixture solution and sonicated for 1 h. Then, the mixture was reacted at 120 °C for 20 h. Next, the solution was neutralized by a sodium hydroxide solution. The final product was further dialyzed for 3 days in a dialysis bag (retained molecular weight: 2000 Da) for purification. The size and shape of GQDs were characterized by TEM (JEOL 2011). The chemical composition was performed by FTIR spectroscopy (Jasco FTIR-6300 spectrometer). The light emission property of GQDs was measured by a plate reader (Tecan infinite 200Pro). The zeta potential was measured by Malvern Zetasizer Nano ZS. The absorption of GQDs was tested by CD spectroscopy (Jasco J-1700 Spectrometer).

### Synthesis of Cys-GQDs and EDA-GQDs

The synthesis of Cys-GQDs and EDA-GQDs was carried out using 1-ethyl-3-(3-dimethylaminopropyl) carbodiimide (EDC) / N-hydroxysuccinimide (NHS) coupling reaction.^36, 51^ 1 mL of EDC with a concentration of 100 mM was mixed with 25 mL GQDs (0.25 mg/mL). After 30-min stirring, 1 mL of NHS (500 mM) was added into the mixture solution and stirred for 30 min. For Cys-GQDs, 1 mL of D-cysteine (100 mM) was added into the reaction and the mixture was reacted for 16 h. For EDA-GQDs, 0.78 mL of EDA was added into the solution and the mixture was reacted for 48 h. The product was purified by a dialysis bag with 1K MWCO.

### Tau_P301L_ expression and purification

Human tau_P301L_ (0N4R) with an N-terminal His_6_ tag was purified following the previous protocol with a slight modification.^52^ Briefly, transformed BL21 (DE3) cells were grown in LB + Kanamycin media at 37°C until OD_600_ reached between 0.6 - 0.8 and were then induced with 0.5 mM IPTG overnight at 16 °C. Cells were then harvested, resuspended, and lysed by probe sonication in the lysis buffer containing 20 mM Tris, 500 mM NaCl, 10 mM imidazole, and 5 mM serine protease inhibitor PMSF, adjusted to pH 8.0. The lysate was then boiled for 20 minutes in a water bath and the debris was pelleted by centrifugation at 20,000g for about 40 min at 4 °C. The supernatant obtained was then injected into a 5 mL IMAC Ni-Charged affinity column and eluted over a gradient of 10–200 mM imidazole. Eluted tau-containing fractions were further purified using GE HiPrep 16/60 Sephacryl S-200 high-resolution size exclusion chromatography into a storage buffer containing 20 mM Tris, 150 mM NaCl, and 1 mM DTT, adjusted to pH 7.6. The purity of the protein was confirmed by sodium dodecyl sulfate-polyacrylamide gel electrophoresis (SDS-PAGE) analysis, and the concentration was estimated using bicinchoninic acid (BCA) assay.

### ThT aggregation assay

Recombinant tau_P301L_ (10 μM final concentration) and GQDs, Cys-GQDs, or EDA-GQDs (final concentration: 6.25, 3.125, 1.25, 0.625, and 0.3125 μM) were mixed in an aggregation buffer (100 mM sodium acetate, 10 μM ThT, 10 μM heparin and 2 mM DTT, pH: 7.4). 200 μL of the same solution was added into two wells of a 96-well plate for each sample. The plate was then sealed with a clear sealing film and allowed to incubate at 37 °C in a Tecan infinite 200Pro plate reader. An automated method was used to carry out ThT fluorescence measurements after 30 s shaking at an excitation wavelength of 450 nm and an emission wavelength of 485 nm at an interval of every 5 min for 45 h. Every experiment included control wells that lacked tau_P301L_, heparin, or GQDs. The aggregation curves were fitted to a Gompertz growth function according to

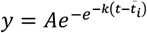

where ***y*** is defined as the normalized ThT fluorescence at time ***t, t***_***i***_ is the inflection point corresponding to the time of maximum growth rate, ***A*** is the maximum normalized ThT fluorescence for a given sample, and ***k*** is the apparent elongation rate constant (***k***_***app***_), in units of h^-1^. The lag time of the aggregation curve can be calculated by

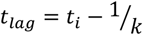

IC_50_ was obtained by fitting the dose-response results using a four-parameter logistic regression model, shown in the following equation.

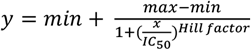

where ***y*** is defined as the endpoint normalized ThT fluorescence at concentration ***x, hill factor*** is the unitless slope factor, and ***max*** and ***min*** correspond to the maximum and minimum endpoint normalized ThT fluorescence in the range of measured concentration, respectively.

### Circular dichroism spectrometry

CD spectra were measured by CD spectroscopy (Jasco J-1700 Spectrometer) at 20 °C. 10 μM of recombinant tau_P301L_ and 6.25 μM of GQDs, Cys-GQDs, or EDA-GQDs were mixed in an aggregation buffer (100 mM sodium acetate, 10 μM ThT, 10 μM heparin and 2 mM DTT, pH: 7.4) and incubated for 4 days at 37 °C. Then, the tau mixture solution was diluted to 0.5 μM for measurement. Spectra were scanned from 190 nm to 260 nm, at 0.1 nm intervals, 5 nm bandwidth, and a scan speed of 50 nm/min.

### Transmission electron microscopy

The preparation procedure of tau aggregation for TEM was the same as CD measurement above. A 3 μL droplet of the tau aggregation solution (10 μM) was placed on the carbon-coated copper TEM grid and wicked using filter paper after 45 s. Then, the grid was washed with 5 μL distilled water twice, stained with 5 μL of 2% (w/v) uranyl acetate for 45 s. Grids were blotted and air-dried before imaging. A JEOL 2011 TEM was used for imaging at an accelerating voltage of 120 kV. Using ImageJ software, the lengths of amyloid fibrils were measured from the average of 16 TEM images per sample.

### ThT disaggregation assay

The tau_P301L_ was diluted to a final concentration of 10 μM in an aggregation buffer (100 mM sodium acetate, 10 μM heparin, 2 mM DTT, and 10 μM ThT, pH: 7.4). The protein was incubated in a microcentrifuge tube for 4 days at 37 °C. After incubation, 200 μL of the same mixture solution (5 μM tau fibers and 6.25 μM GQDs, Cys-GQDs, or EDA-GQDs) was added into two wells of a 96-well plate for each sample. The plate was then sealed with a clear sealing film and allowed to incubate at 37 °C in a Tecan infinite 200Pro plate reader. An automated method was used to carry out ThT fluorescence measurements after 30 s shaking at an excitation wavelength of 450 nm and an emission wavelength of 485 nm at an interval of every 5 min for 30 h. Every experiment included control wells that lacked tau_P301L_ fibrils or GQDs.

### Cellular seeding assay

HEK293 cells stably expressing tau-RD (LM)-YFP were cultured in DMEM complete medium containing 10% FBS, 1% penicillin/streptomycin, and 1% Glutamax under 5% CO_2_ at 37 °C. 90 μL of cells were plated at a density of 15,000 cells/well in a 96-well tissue culture plate.

For seeding by monomeric tau, 10 μM of tau_P301L_ was incubated in an aggregation buffer containing 100 mM sodium acetate, 10 μM heparin, 2 mM DTT, and GQDs or Cys-GQDs (final concentration: 6.25 and 31.25 μM), pH: 7.4 for 4 days at 37 °C. Control vials included those in which buffer was added in place of tau_P301L_ or heparin. Following tau incubation, 8 μL of tau aggregation solution was mixed with 32 μL of low-serum Opti-MEM medium and 2 μL of Lipofectamine 2000, and the mixture solution was incubated for an additional 20 min at room temperature.

For seeding by fibrillar tau, the tau_P301L_ was diluted to a final concentration of 10 μM in an aggregation buffer containing 100 mM sodium acetate, 10 μM heparin, and 2 mM DTT, pH: 7.4. The protein was incubated for 4 days at 37 °C. Control vials included those in which buffer was added in place of tau_P301L_ or heparin. Following incubation, 8 μL of tau fibrils were mixed with 28 μL of low-serum Opti-MEM medium and 4 μL of GQDs or Cys-GQDs with different concentrations. The mixture was incubated at 37 °C for 36 h, mixed with 2 μL of Lipofectamine 2000, and incubated for 20 min at room temperature.

Then, 10 μL of mixture solution was added to HEK-293 cells and incubated for 48 h at 37 °C. Every experiment included control wells that lacked tau_P301L_, heparin, or GQDs. After that, cells were detected by a BioTek Cytation 5 cell imager and a microplate reader. 10 × 10 pictures/well were taken at 20× magnification under a fluorescein isothiocyanate (FITC) channel (ex: 469 nm/em: 525 nm), and the punctate counting was carried out using built-in software.

### Fluorescence quenching assay

10 μM of tau_301L_ in 100 mM sodium acetate buffer was incubated with different concentrations of GQDs or Cys-GQDs (6.25, 12.5, and 25 μM) at room temperature in a 96-well plate. After 10-min incubation, the fluorescence of the tau mixture solution was detected at an excitation wavelength of 265 nm and an emission wavelength from 295 nm to 380 nm using a Tecan infinite 200Pro plate reader. All spectra were subtracted with GQD background solutions.

### CCK-8 assay

SH-SY5Y human neuroblastoma cells were cultured in the DMEM/F12 complete medium containing 10% FBS and 1% penicillin/streptomycin under 5% CO_2_ at 37 °C. 100 μL of cells were plated at a density of 10,000 cells/well in 96-well tissue culture plates and incubated overnight. Control wells contained only medium for background measurement. Then, add 10 μL of GQDs with a concentration of 15 μM and 50 μM into cells, and add 10 μL of PBS as the negative control. Incubate the 96-well microplates for 24 or 48 h in a CO_2_ incubator at 37 °C. After that, change the medium, add 10 μL of Cell Counting Kit-8 reagent to cells and incubate the 96-well microplates for 3 h at 37 °C. The absorbance was measured at 450 nm using a Tecan infinite 200Pro plate reader. The cell viability was calculated using the following equation.

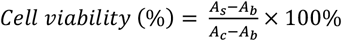

where ***A***_***S***_ and ***A***_***C***_ are the absorbances measured at 450 nm for cells added GQDs and PBS, respectively, and ***A***_***b***_ is the absorbance measured at 450 nm for only medium.

## Conclusions

The aggregation of tau proteins plays a key role in the observed clinical symptoms and pathology of tauopathies. However, current drug candidates for the treatment of tauopathy have not been very successful in clinical trials. Therefore, there is an urgent need for novel modulators of tau fibrillization and propagation. Here, we engineered the size and charge of GQDs and investigated their interaction with tau. *In vitro* and cellular assays suggest that GQDs with larger size and negative charge can inhibit the aggregation of tau more efficiently *via* electrostatic interactions and π-π stacking with tau. In addition, GQDs with positive charge are more effective in the disassembly of mature tau fibrils. Given that GQDs can cross the BBB and be conjugated to targeting ligands, our results contribute to the future development of GQD-based drug delivery systems for the treatment of tauopathies.

## Author Contributions

Y.W. and J.R.D. originated and conceptualized the study. R.Z. designed and carried out the experiments. Y.Z. developed the methodology of synthesis and functionalization of GQDs. K.M.M. and B.H.R. expressed and purified tau_P301L_. K.M.M. developed the methodology of tau biosensor cellular seeding assay. R.Z. applied statistical and mathematical techniques to analyze data. Y.W. and J.R.D. supervised all the work. R. Z. prepared the manuscript, and all authors provided input on data interpretation, discussions, and writing.

## Acknowledgements

We acknowledge funding from an American Cancer Society Institutional Research Grant (ACS IRG-17-182-04 to Y.W.) and funding to Y.W. from the National Science Foundation Industry-University Cooperative Research Center (The Center for Bioanalytic Metrology). This work was also supported by a grant from the National Institutes of Health (R01AG074570 to J.R.D.) and an Indiana Clinical and Translational Sciences Institute TL1 predoctoral fellowship (to B.R.). TEM images were carried out in part in the Integrated Imaging Facility, University of Notre Dame, using JEOL 2011 TEM. We thank Maksym Zhukovskyi for the knowledge and expertise as well as time towards this research. The authors would like to thank James Johnston, Hyunsu Jeon, and Gaeun Kim for the helpful discussion on the results and experiments. The authors are also grateful to Syrah Starnes and Isaac Angera for their assistance with experiments. Fig. 2b, 4a, 5a, and 8 were created using BioRender.com and used with permission.

