## Supporting Information for "Inhibition and Disassembly of Tau Aggregates by Engineered Graphene Quantum Dots"

### Supporting Figures

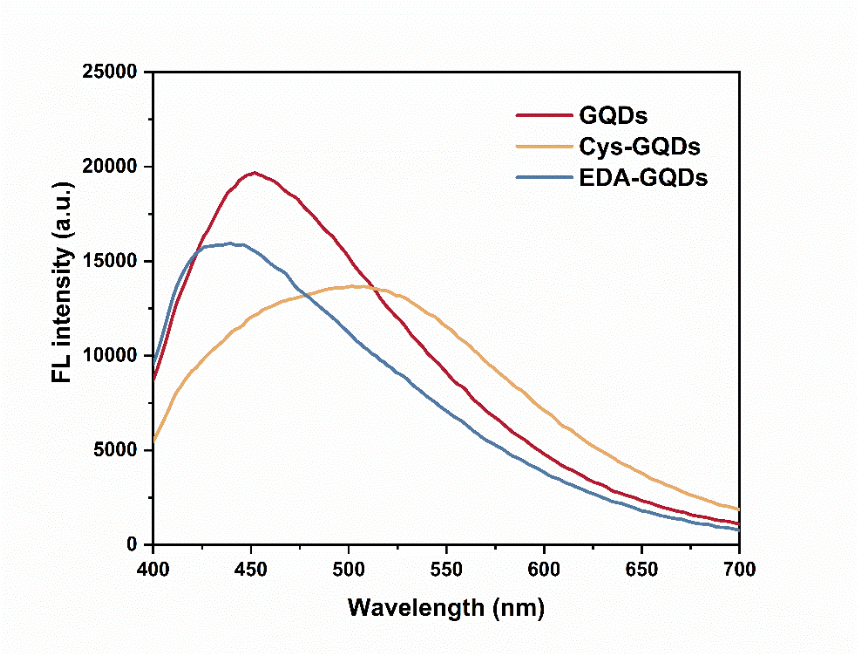


**Fig. S1** Fluorescence spectra of GQD, Cys-GQDs, and EDA-GQDs excited at 365 nm.

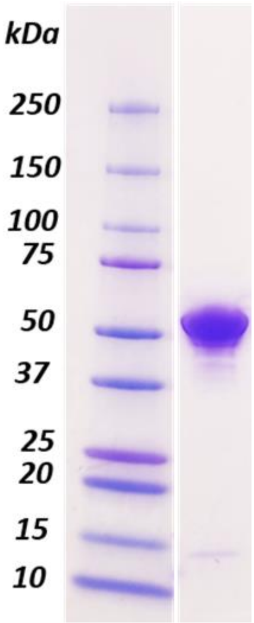


**Fig. S2** SDS/PAGE (Coomassie blue stain) of final purified $tau_{P301L}$ protein.

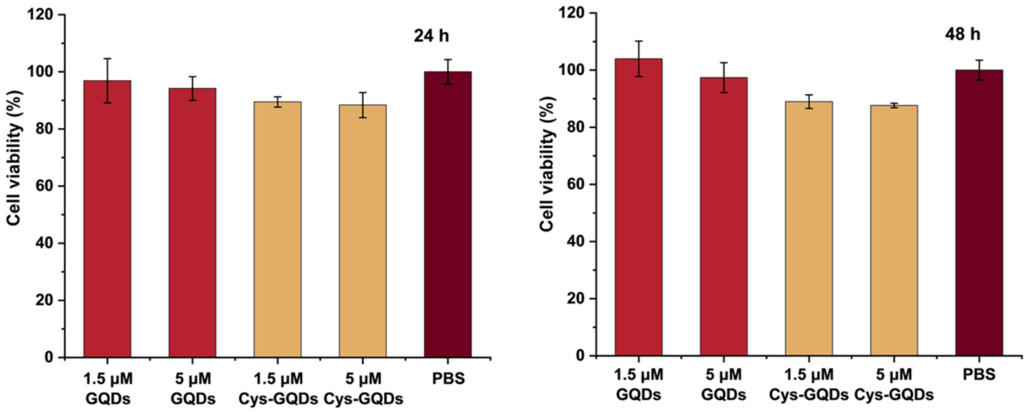


**Fig. S3** The cell viability of SH-SY5Y human neuroblastoma cells after incubation with 1.5 μM and 5 μM of GQDs and Cys-GQDs for 24 and 48 h, tested by CCK-8 assay.

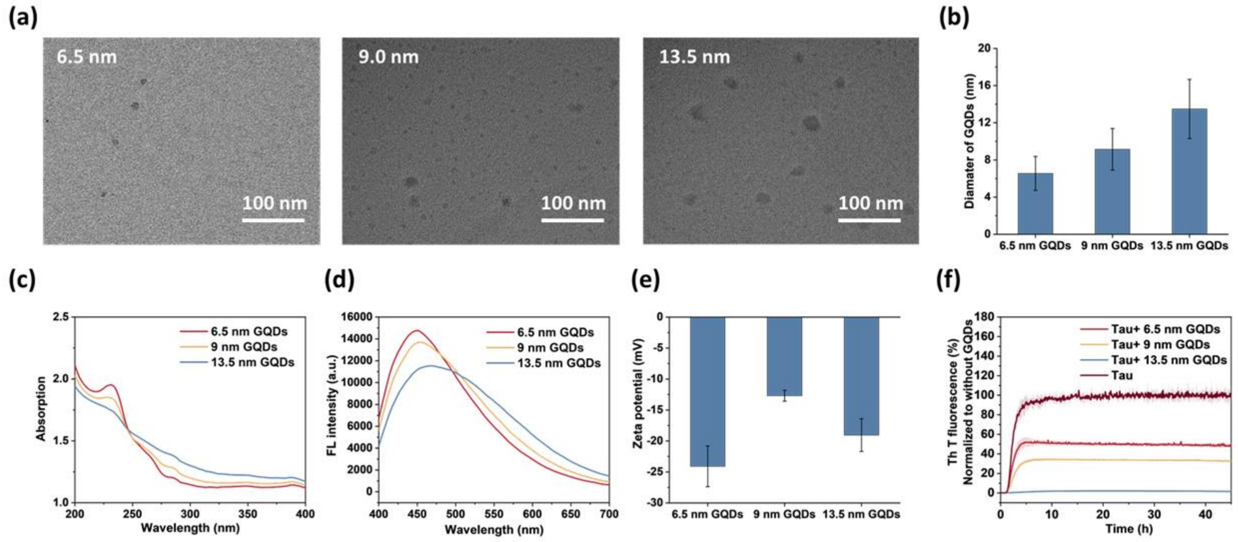


**Fig. S4** The size effect of GQDs on the tau aggregation. (a) GQDs were separated using centrifuge filters with 3 KDa and 10 KDa molecular weight cutoffs. TEM images of GQDs with average diameters of 6.5, 9.0, and 13.5 nm. (Scale bar: 100 nm) (b) Size distribution of GQDs with average diameters of 6.5 nm, 9.0 nm, and 13.5 nm. (c) Absorption spectra, (d) fluorescence spectra (excited at 365 nm), and (e) zeta potentials of 1.25 μM GQDs with three different diameters. (f) $Tau_{P301L}$ aggregation was inhibited with GQDs of different sizes with a concentration of 1.25 μM monitored by ThT fluorescence assay.

### Supporting Tables

**Table S1** The apparent elongation rate constants ($\boldsymbol{k_{app}}$) and lag times ($\boldsymbol{t_{lag}}$) of tau aggregation incubated with/without 6.25 μM of GQDs or functionalized GQDs.

| | Tau aggregation (%) | $\boldsymbol{k_{app}}$ ($h^{-1}$) | $\boldsymbol{t_{lag}}$ (h) |
|---|---|---|---|
| Tau | 100 ± 9.27 | 0.76 ± 0.008 | 1.84 ± 0.011 |
| Tau + GQDs | 2.20 ± 0.23 | 0.70 ± 0.039 | 3.46 ± 0.063 |
| Tau + Cys-GQDs | 2.01 ± 0.082 | 0.38 ± 0.021 | 3.88 ± 0.096 |
| Tau + EDA-GQDs | 11.2 ± 0.10 | 0.95 ± 0.034 | 0.95 ± 0.030 |

**Table S2** Average length and density of tau fibers were obtained from TEM images. The length distribution is close to an exponential distribution, and thus its standard deviation is close to the mean value.

| | Average length of tau fibers (nm) | Average density (fibers/μ$m^2$) | Average length of tau fibers in the unit area (nm/μ$m^2$) |
|---|---|---|---|
| Tau+ GQDs | 277 ± 302 | 1.27 ± 0.48 | 371 |
| Tau+ Cys-GQDs | 333 ± 328 | 1.66 ± 0.98 | 506 |
| Tau+ EDA-GQDs | 571 ± 707 | 3.76 ± 0.84 | 2033 |
| Tau | 1623 ± 1255 | 4.53 ± 2.08 | 7352 |
